# Does 10-Hz cathodal oscillating current of the parieto-occipital lobe modulate target detection?

**DOI:** 10.1101/235416

**Authors:** Sarah S. Sheldon, Kyle E. Mathewson

**Affiliations:** Department of Psychology, Faculty of Science, University of Alberta, Edmonton, Alberta, Canada; Neuroscience and Mental Health Institute, Faculty of Medicine and Dentistry, University of Alberta, Edmonton, Alberta, Canada

**Keywords:** alpha oscillations, transcranial current stimulation, entrainment, detection, negative findings

## Abstract

The phase of alpha (8-12 Hz) brain oscillations have been associated with moment to moment changes in visual attention and awareness. Previous work has demonstrated that endogenous oscillations and subsequent behavior can be modulated by oscillating transcranial current stimulation (otCS). The purpose of the current study is to establish the efficacy of cathodal otCS for modulation of the ongoing alpha brain oscillations, allowing for modulation of individual’s visual perception. Thirty-six participants performed a target detection with sham and 10-Hz cathodal otCS. Each participant had two practice and two experimental sets composed of three blocks of 128 trials per block. Stimulating electrodes were small square sponges (20 cm^2^) placed on the participant’s head with the anode electrode at Cz and the cathode electrode at Oz. A 0.5 mA current was applied at the cathode electrode every 100 ms (10 Hz frequency) during the otCS condition. The same current and frequency was applied for the first 10-20 s of the sham condition, after which the current was turned off. Target detection rates were separated into ten 10-ms bins based on the latency between the stimulation/sham pulse and target onset. Target detection rates were then compared between the sham and otCS experimental conditions across the ten bins in order to test for effects of otCS phase on target detection. We found no significant difference in target detection rates between the sham and otCS conditions, and discuss potential reasons for the apparent inability of cathodal otCS to effectively modulate visual perception.

## 1 INTRODUCTION

Oscillating neural activity enables the brain to communicate and coordinate across different areas in order to carry out important cognitive functions. Over the last decade, there has been a resurgence of interest in oscillatory activity due to recent technological advances that enable non-invasive modulation of these brain oscillations (Fröhlich, 2015; Fröhlich et al., 2015; Herrmann et al., 2013). In particular, transcranial current stimulation (tCS) has become a popular method because it provides the possibility to modulate the phase, amplitude, and frequency of ongoing oscillatory activity (Paulus, 2011).

The most common applications of tCS involves the delivery of the electrical stimulation as either a direct current (i.e., current of a constant intensity and polarity) or an alternating current (i.e., current that oscillates between negative and positive polarity). Anodal (positive polarity) and cathodal (negative polarity) transcranial direct current stimulation (tDCS) can modulate the neuronal response threshold by inducing depolarization or hyperpolarization, respectively (Jackson et al., 2016; Paulus et al., 2016). On the other hand, transcranial alternating current stimulation (tACS) can modulate ongoing neuronal activity in a frequency-specific manner. It is thought that tACS effects response thresholds in a manner similar to tDCS except that alternating between positive (anodal) and negative (cathodal) current results in the neural oscillations becoming entrained to the timing of the alternating current (Antal and Paulus, 2013; Herrmann et al., 2013; Reato et al., 2013; Vosskuhl et al., 2015). This means that tACS can be used to manipulate oscillatory activity in an experimental setting to understand the relevance of such induced oscillations for cognition.

Although numerous studies have demonstrated that these electrical stimulation methods can effect perception (Antal et al., 2004; Antal and Paulus, 2008; Helfrich et al., 2014; Neuling et al., 2012a) and cognition (Antonenko et al., 2016; Marshall et al., 2006; Simonsmeier and Grabner, 2017; Zaehle et al., 2011), it seems that many other studies have found little to no evidence supporting the efficacy of these techniques (Brignani et al., 2013; Horvath et al., 2015; Marshall et al., 2016; Tremblay et al., 2014; Veniero et al., 2017). Therefore, here we used a well studies paradigm of alpha oscillations affecting visual perception as a test of the feasibility of using tACS to manipulate oscillations and cognition.

Using tACS with a DC-offset is referred to as oscillating transcranial current stimulation (otCS). This technique can be thought of as a combination of tDCS and tACS., and this combination of tDCS and tACS has been shown to be effective for boosting memory (Marshall et al., 2006), and pulsed current stimulation has been shown to affect corticospinal excitability (Jaberzadeh et al., 2014). We therefore utilized otCS here to manipulate posterior parietal alpha oscillations and test if there was any influence on target detection.

Brain oscillations within the alpha (8-12 Hz) frequency band have emerged as a marker of visual perception and selective attention (Mathewson et al., 2011). We and others have shown that target detection depends on the phase of alpha oscillations at the moment of target onset (Mathewson et al., 2009), which we have explained due to alpha acting as a pulsating inhibition in the brain. We have found using fast optical imaging that these alpha oscillations relevant for detection can be localized to the posterior parietal cortex (Mathewson et al., 2014). We have found support for this theory in a series of studies in which we rhythmically entrain alpha oscillations with visual stimulation and observe subsequent rhythmic modulation in target detectability (Kizuk and Mathewson, 2017; Mathewson et al., 2012, 2014). We find that 12-Hz rhythmic visual stimulation induces phase locking at the same frequency in the EEG, as well as these fluctuations in target detection. In comparison to the classical rhythmic sensory stimulation protocols which entrain the entire visual system, the use of tCS offers the advantage of directly stimulating cortical targets (Brignani et al., 2013).

The aim of the current study was to provide a proof of principle that the entrainment of ongoing neural oscillations by rhythmic visual stimulation can be replicated with cathodal otCS at the same frequency. The present study aims to address this issue by attempting to control the phase alpha oscillations in the posterior parietal cortex during visual perception. We chose otCS because it has been associated with modulation of parieto-occipital alpha activity and subsequent behavior (Kasten and Herrmann, 2017).

## 2 MATERIALS AND METHODS

### 2.1 Participants

Thirty-six participants from the University of Alberta community participated in the study (mean age = 21; age range = 17-32, 10 males). Participants were all right-handed, and had normal or corrected normal vision and no history of neurological problems. All participants gave informed written consent, were either compensated at a rate of $10/hr or given research credit for their time, whichever was applicable. The study adhered to the tenets of the Declaration of Helsinki and was approved by the Internal Ethics Board at the University of Alberta.

### 2.2 Target Detection Task

Participants were seated 57 cm away from a 1920 x 1090 pixel^2^ ViewPixx/EEG LCD monitor (VPixx Technologies, Quebec, Canada) with a refresh rate of 120 Hz, simulating a CRT display with LED backlight rastering. The rastering, along with 8-bit digital TTL output triggers yoked to the onset and value of the top left pixel, allowed for submillisecond accuracy in pixel illumination times, which were confirmed with a photocell prior to the experiment. Stimuli were presented on a 50% gray background using a Windows 7 PC running MATLAB R2012b with the Psychophysics toolbox (Version 3; Brainard, 1997; Pelli, 1997). See Figure 1A for the stimulus dimensions. Video output was sent to the ViewPixx/EEG with an Asus Striker GTX760 (Fremont, CA) graphics processing unit.

**Figure 1.**
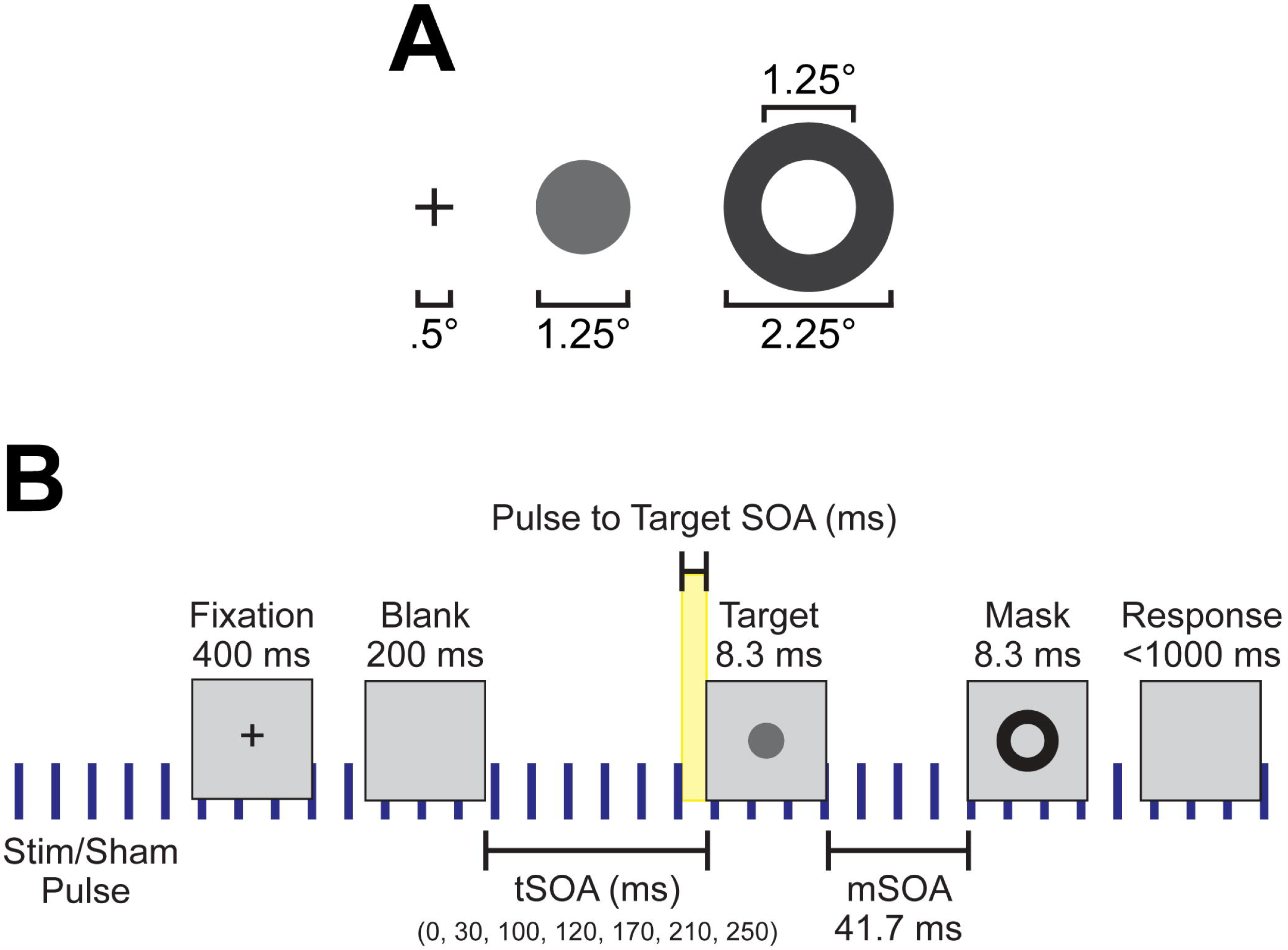
Experimental setup and design. **(A)** Spatial dimensions of the stimuli, which were presented to subjects at the center of the screen. **(B)** Individual trial timeline with durations of each screen presentation. Blue vertical lines indicate the continuous application of the 10 Hz sham or otCS stimulation pulse throughout the task. Highlighted yellow area was the time range between the preceding stimulation (sham or otCS) pulse and the onset of the target which was used to subdivide the trials into 10 ms bins.

Each trial began with a black fixation cross presented at the center of the monitor for 400 ms. The fixation cross was followed by a blank screen. The blank screen remained for 200, 230, 300, 320, 370, 410, or 450 ms (target stimulus onset asynchrony; tSOA) after which the target appeared for 8.33 ms (one monitor refresh). The target was followed by a backward mask lasting for 8.33 ms with a constant 41.7 ms target-mask SOA (mSOA). Following the mask offset, the participant had 1000 ms to respond before the next trial began. There were 128 trials per block, and three blocks per experimental condition. On 20% of trials, the target was omitted to assess false alarms. A summary of the task sequence can be seen in Figure 1B.

In the first two conditions, the target luminance value was adjusted throughout the task based on a 3-up/1-down staircasing procedure that was chosen because it targeted a 0.5 target detection rate for each individual (Garcia-Pérez, 1998; Kingdom and Prins, 2016). The target luminance value in the final two conditions remained constant and determined for each participant by taking the average target luminance value across the last two blocks of trials in the second staircasing block.

### 2.3 Electrical Stimulation

A battery-driven stimulator (Oasis Pro, Mind Alive, Canada) was used to deliver a 10-Hz oscillating cathodal transcranial electrical current via rubber electrodes encased in sponges (5×4 cm; Oasis Pro, Mind Alive, Canada) and soaked in saline solution. The electrodes were attached to the head underneath an EEG Recording Cap (EASYCAP, Herrsching, Germany) with the cathodal electrode (where the current was applied) at Oz and the anodal electrode placed at Cz. These positions were chosen for maximal stimulation intensity in the parieto-occipital cortex (Neuling et al., 2012b). The stimulation current had a rounded square waveform that was delivered at a 10-Hz frequency. The onset of each stimulation pulse was recorded by the amplifier via a customized trigger output added to the Oasis Pro stimulator by the manufacturer with the accuracy confirmed with oscilloscopes prior to the experiment.

The intensity of the stimulation current was adjusted for each participant to ensure that they did not experience pain, tingling or other unpleasant sensations. To obtain this threshold, we started with an intensity level of 0.50 mA (peak-to-peak). If the participant indicated unpleasant sensations, we decreased the intensity in steps of 0.02 mA until the participant reported little to no skin sensation. The obtained threshold level ranged between 0.34-0.50 mA (*M* = 0.46, *SD* = 0.05) was used as stimulation intensity in the tCS condition.

The sham condition consisted of a 10 s fade-in and 20 s of stimulation at 0.50 mA. The current was then shut off by disconnecting the Oasis Pro stimulator from the stimulating electrodes outsight of the sign of the participant. Disconnecting the stimulating device from the electrodes did not interrupt the stimulation triggers sent to the amplifier, which can therefore be used as control timings. The experimental condition also consisted of a 10 s fade-in and 20 s of stimulation at 0.50 mA, after which the current intensity was decreased to the individual’s obtained threshold level.

### 2.4 EEG Recording

During the target detection task, EEG data was recorded using a 16-channel V-amp amplifier (Brain Products, München, Germany) from 15 scalp locations (O1, O2, P7, P3, Pz, P4, P8, T7, C3, Cz, C4, T8, F3, Fz, F4; 10/20 system), a ground electrode at position Fpz, and two reference electrodes, placed at the right and left mastoids, with Ag/AgCl sintered ring electrodes (EASYCAP, Herrsching, Germany) in a 20-channel electrode cap (EASYCAP). SuperVisc electrolyte gel and mild abrasion with a blunted syringe tip were used to lower impedances to less than 5 kΩ for all electrode sites except Cz which did not have direct contact with the head because it was on top of the stimulating electrode sponge. EEG was recorded online referenced to an electrode attached to the left mastoid. Offline, the data were re-referenced to the arithmetically derived average of the left and right mastoid electrode.

In addition to the 15 EEG sensors, two reference electrodes, and the ground electrode, the vertical and horizontal bipolar EOG was recorded from passive Ag/AgCl Easycap disk electrodes affixed above and below the left eye, and 1 cm lateral from the outer canthus of each eye. Prior to placement of electrodes, the participant’s skin was cleaned using Nuprep (an exfoliating cleaning gel) and electrolyte gel was used to lower the impedance of these EOG electrodes to under 5 kΩ in the same manner as previously mentioned. The bipolar vertical and horizontal EOG was recorded using a pair of BIP2AUX converters in the V-amp auxiliary channels (Brain Products). The EOG electrodes had a separate ground electrode affixed to the central forehead.

Data were digitized at 2000 Hz with a resolution of 24 bits (0.049 μV steps). Data were collected inside a sound and radio frequency-attenuated chamber (40A-series; Electro-Medical Instruments, Mississauga, Ontario, Canada), with copper mesh covering a window. The lights were left on, and the window was covered during experiments. The only electrical devices inside the chamber were the amplifier, powered from a battery powered laptop located outside the chamber, speakers, keyboard, and mouse, all powered from outside the room, the ViewPixx monitor, powered with DC power from outside the chamber, and a battery-powered intercom. Nothing was plugged into the internal power outlets, and any electrical devices (e.g., cell phones) were removed from the chamber during recording.

### 2.5 Design and Procedure

For all the participants, the study consisted of one session and took approximately 90 minutes. We implemented a single-blind sham-controlled design in which participants underwent two experimental conditions (otCS and sham) in a counterbalanced order. EEG data was simultaneously recorded during both conditions.

The procedure started with the participants performing three practice blocks of the staircased version of the target detection task while the experimenters set-up the EEG cap and electrical stimulation electrodes. After the practice blocks and set-up, the electrical stimulation intensity was determined for each participant using the procedure described above. Next, the participant performed the staircased version of the target detection task a second time under the stimulation sham condition. The average luminance value of the target in the last two blocks of trials was calculated for each participant. Finally, participants performed the target detection task under the otCS and sham experimental conditions (counterbalanced across subjects) using the previously calculated target luminance value.

Although EEG data was recorded throughout the final three conditions, attempts to remove the otCS stimulation artifact with both traditional and advanced multi-step procedures (Helfrich et al., 2014; Kohli and Casson, 2015; Liu et al., 2012) were unsuccessful. This was most likely due to the presence of small fluctuations of stimulation intensity caused by the stimulating device. Therefore, were not able to examine possible psychophysiological effects.

### 2.6 Questionnaire

To obtain possible adverse effects for otCS, a version of a questionnaire introduced by Brunoni et al. (2011) was used. The following side-effects were inquired: headache, neck pain, scalp pain, tingling, itching, burning sensation, skin redness, sleepiness, trouble concentrating and acute mood change. Participants were asked to indicate the intensity of the side-effect (1, absent; 2, mild; 3, moderate; 4, severe) and if they attributed the side-effect to the tACS (1, none; 2, remote; 3, possible; 4, probable; 5, definite).

The most reported adverse effects (intensities rated higher than 1) after the experiment were trouble concentrating (70.0%), sleepiness (66.7%) and scalp tingling (56.7%). Ratings for intensity of adverse effects were generally relatively low, except for sleepiness (*M* = 2.12) and trouble concentrating (*M* = 2.10). For the ratings on whether subjects attributed the adverse effects to the stimulation, only tingling achieved an average score above 2 (*M* = 2.20).

### 2.7 Data Analyses

Data analysis was performed using MATLAB R2017a (The MathWorks Inc, Natick, MA, USA) and EEGLAB 13.6.5b (Delorme and Makeig, 2004). All statistical analyses were conducted using SPSS 11.5.0 (Chicago, IL) and R 3.3.1 (R Core Team, 2013).

#### 2.7.1 Target detection performance

First, the trials from the non-staircased version of the target detection task were subdivided into 10 ms bins based on the time between the preceding stimulation pulse and the onset of the target (pulse to target SOA; see Figure 1B). This was our main independent variable, since we predict that if alpha oscillations are being entrained by the electrical stimulation their phase should influence detection. Because a stimulation pulse was every 100 ms, this meant that there was a total of ten bins. Target detection rates (proportion of targets participants detected) of each participant was calculated for these ten 10 ms bins after excluding catch trials (where no target appeared) and trials without a valid response. These calculations were performed separately for each stimulation condition (otCS and sham). A test of the mean detection rates across bins between otCS and sham conditions was conducted using a mixed ANOVA where the 10 ms bins and stimulation condition were within-subject factors, condition order (otCS before sham or sham before otCS) was a between-subject factor, and the participants were treated as a random variable. The ANOVA was performed in R using the built-in aov function and the ezANOVA function from the ez package (Lawrence, 2016). The analysis yielded a significant interaction between stimulation condition and order of conditions indicating the presence of a sequence effect (see Results section and Figure 3A). The sequence effect was not relevant to the hypothesis that target detection rates will vary in a sinusoidal manner relative to otCS stimulation pulses but not the sham pulses. Therefore, the target detection rates were normalized for each participant in each condition separately and then re-tested with the mixed ANOVA.

Finally, the behavioral data was subdivided into twelve bins of 32 consecutive trials across the three blocks of each stimulation condition and submitted to a repeated-measures ANOVA. This was done to investigate whether there was a change in target detection rates across the condition, since if alpha power increases with stimulation time target detection should get worse.

#### 2.7.2 Sinusoidal model of detection rates

For each participant and stimulation condition, the sinusoidal function

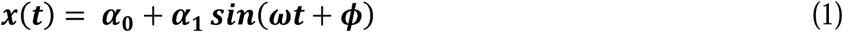

with intercept α_0_, amplitude α_1_, and phase ϕ was estimated for the standardized target detection rates of the ten 10-ms bins in each stimulation condition. The routine to fit the parameters was initialized with random start values, and used a nonlinear least-squares method. The parameters were limited by the following constraints: ϕ ∈ (–π, π); α_1_ ∈ (0,∞); and, frequency ω was fixed at 0.06 bins/cycle (100 Hz). To compare the influence of the otCS and sham stimulation pulses on target detection rates, a paired Student’s *t* test was performed on the estimated amplitude (α_1_) and a Wilcoxon signed-ranks test was performed on the goodness-of-fit measure adjusted r-square 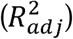.

#### 2.7.3 EEG data

The average voltage in the 300 ms baseline prior to the target was subtracted on each trial for every electrode. Trials with absolute voltage fluctuations on any channel greater than 1000 μV were discarded, and data was segmented into 1800 ms epochs aligned to target onset (-800 ms pre-target onset to 1000 ms post-target onset). Eye movements were then corrected with a regression-based procedure developed by Gratton, Coles, and Donchin (1983). After a second baseline subtraction with 300 ms pre-target, trials with remaining absolute voltage fluctuations on any channel greater than 500 μV were removed from further analysis.

## 3 RESULTS

The mixed ANOVA on the mean detection data yielded no significant main effects or interactions (Figure 2A) except for the interaction between stimulation condition and stimulation condition order (*F*(1,646) = 38.20, *p* < 0.001). This indicates that there was a sequence effect in that mean target detection rates were greater in the second stimulation condition compared to the first, regardless of whether sham came before otCS (sham condition: *M* = 0.46, *SE* = 0.05; otCS condition: *M* = 0.51, *SE* = 0.04) or otCS came before sham (sham condition: *M* = 0.49, *SE* = 0.05; otCS condition: *M* = 0.45, *SE* = 0.04). All other main effects and interactions had an *F*-value of less than 1.

**Figure 2.**
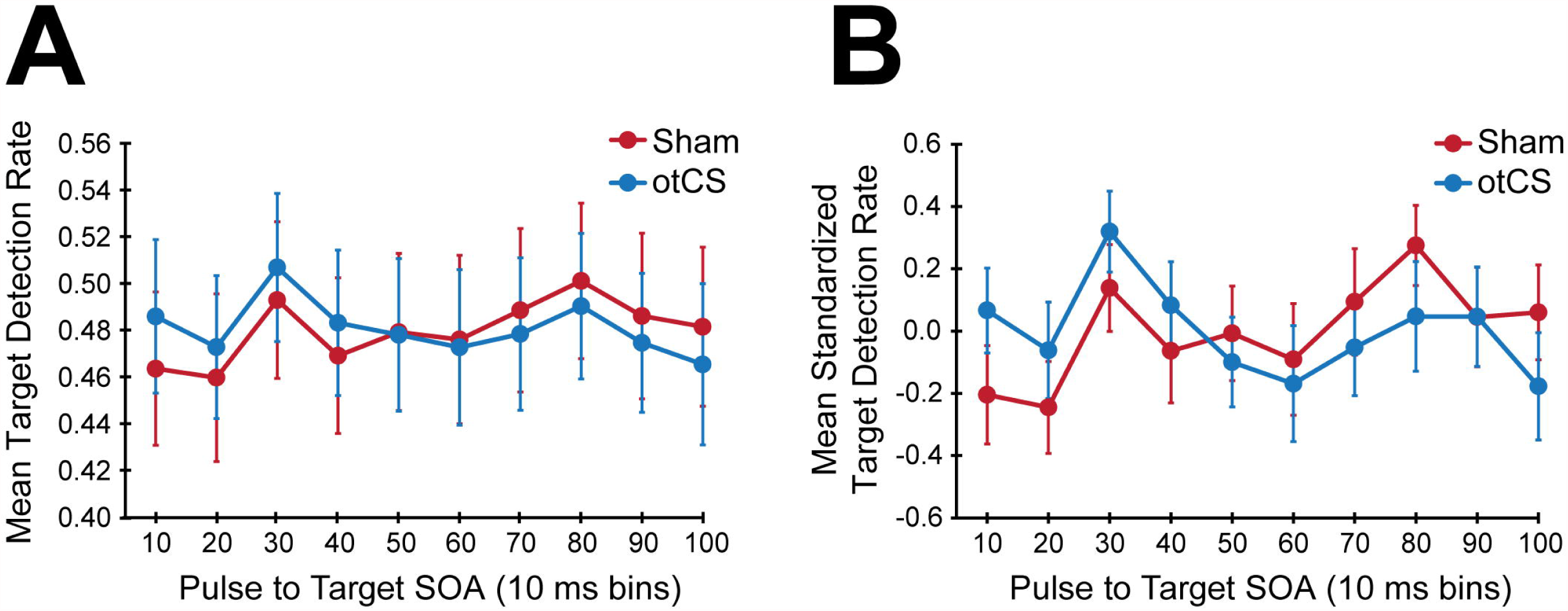
**(A)** Mean target detection rates and **(B)** mean standardized target detection rate in each 10 ms bin during the sham and otCS stimulation conditions. Error bars indicate the standard error (*SE*).

**Figure 3.**
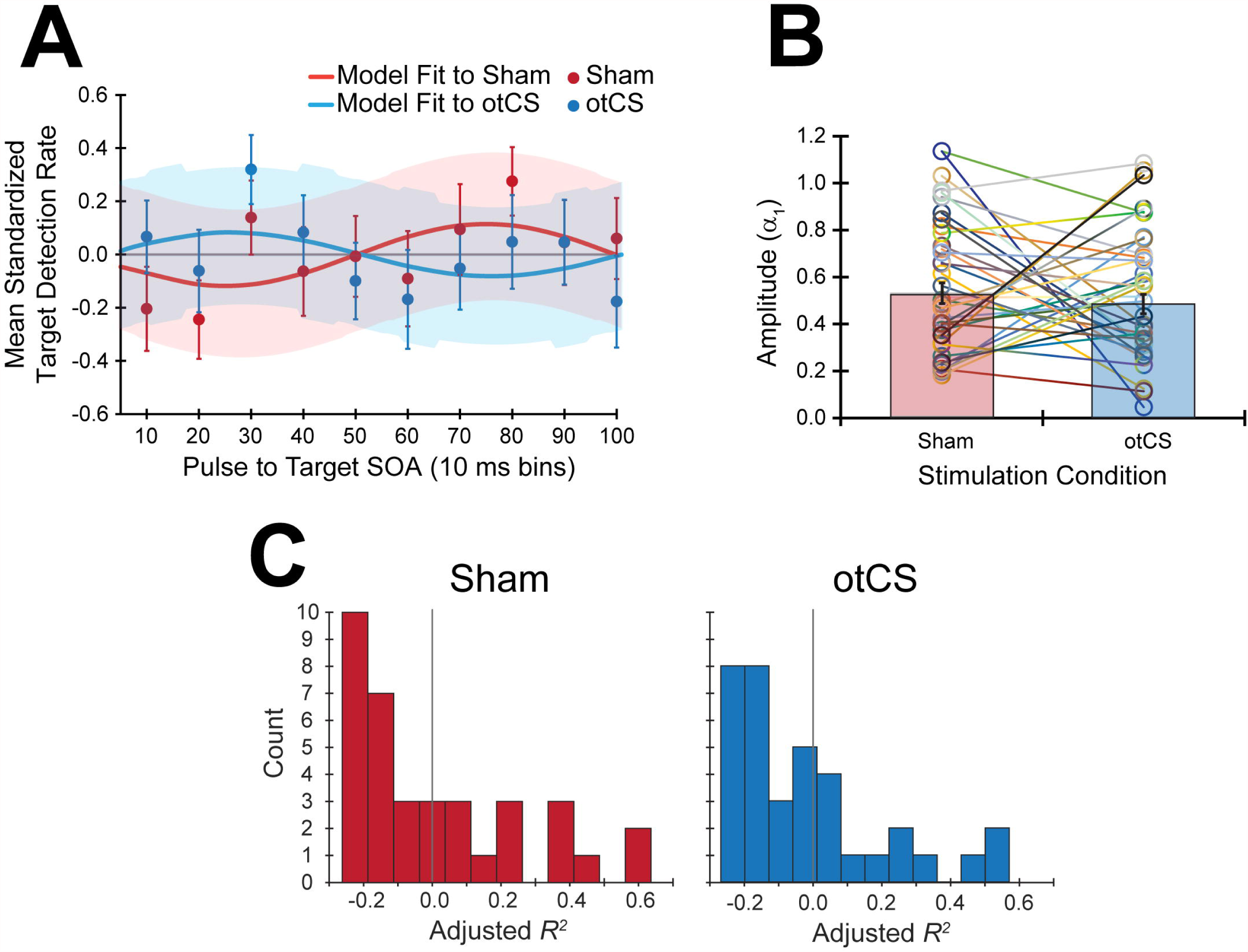
**(A)** The mean standardized target detection rates in each 10 ms bin for the sham and otCS stimulation conditions overlaid by each fitted sine functions for the sham and cathodal otCS stimulation conditions. Error bars and shaded color regions indicate the *SE* of the mean standardized detection rates and model fits, respectively. **(B)** The open circles denote the individual amplitude (α_1_) estimates for each participant in the sham and otCS conditions. Lines connect the data points from the same participant. The red and blue bars are the group averages in the sham and otCS conditions, respectively. The error bars are the *SE*. (C) Histograms of the goodness-of-fit measure, adjusted *R*^2^, of the sinusoidal model to the mean standardized target detection rates in sham (left) and otCS (right) stimulation conditions. The larger the adjusted r-square value, the more variability in the detection rates explained by the model. Grey line marks a value of zero.

To compensate for this sequence effects, target detection rates were normalized for each participant in each condition separately and were tested again with the same ANOVA. The statistical test also yielded no significant main effects or interactions including the interaction between stimulation condition and stimulation condition order (Figure 2B). There were no main effects or interactions with a *p*-value less than 0.20.

Contrary to our hypothesis, the sinusoidal pattern of the target detection rates did not seem to be strongly modulated by the cathodal otCS stimulation pulses compared to the sham (Figure 3A). This is supported by a paired *t* test which indicates that there is no significant difference in the estimated amplitude parameters (α_1_) from the fitted sine functions to the otCS and sham behavioral data (*t*(35) = 0.65, *p* = 0.52; Figure 3B). Furthermore, a Wilcoxon signed-ranks test indicated that the amount of variability in the target detection rates accounted for by the sinusoidal model (adjusted *R^2^* value) did not differ significantly between the sham and otCS stimulation conditions (*Z* = -0.58, *p* = 0.56; Figure 3C).

Finally, the mean target detection rates across each experimental condition was examined to see if there was an effect of the otCS stimulation over the course of the trials. Mauchly’s test indicated that the assumption of sphericity was violated for the stimulation condition x bins interaction, *W* = 0.065, p < .01, ε = .66. The degrees of freedom were corrected using Greenhouse-Geisser estimates of sphericity. There was a significant main of bin on target detection rates (*F*(11,385) = 4.78, *p* < 0.001). There was no significant main effect of stimulation condition (*F*(1,35) < 1.00), nor a significant interaction between stimulation condition x bins (*F*(7.23,253.19) = 0.65, *p* = 0.72). As can be seen in Figure 4, there was a change in target detection rates across the duration of the task, but this change was about the same in both conditions. A post hoc test using the Holm procedure to control for Type I errors revealed that the first 32 trials (bin 1; *M* = 0.57, *SE* = 0.03) had significantly better target detection rates than the set of trials in bin 4 (*M* = 0.46, *SE* = 0.03), bin 7 (*M* = 0.47, *SE* = 0.02), and bin 8 (*M* = 0.48, *SE* = 0.02). Because participants performed the task in three blocks of 128 trials, the end of the first block corresponds to bin 4 and the end of the second block corresponds to bin 8. Therefore, the most likely explanation for these results is that the participants got fatigued towards the end of each block.

**Figure 4.**
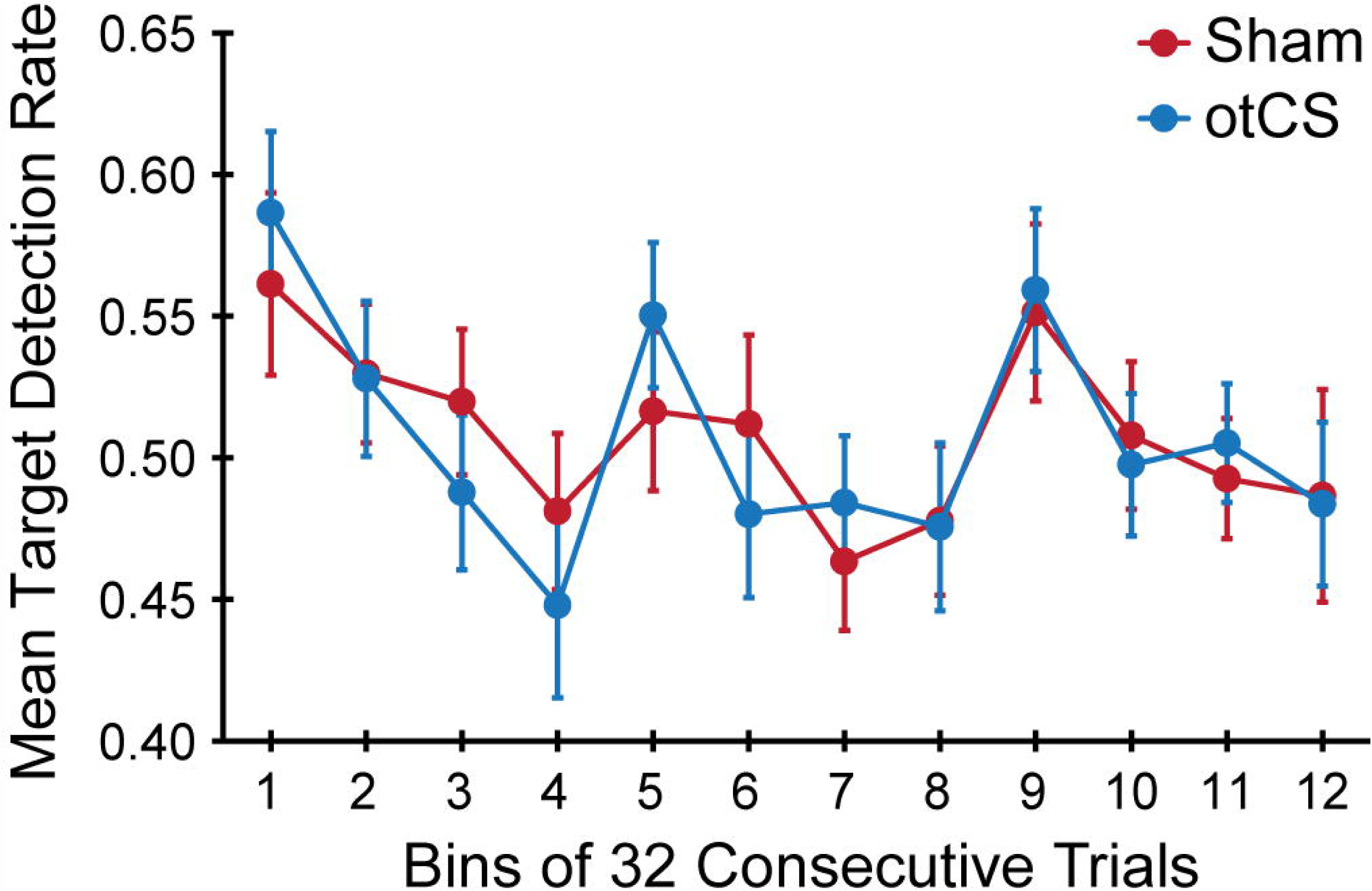
Mean target detection rates rate in each bin of 32 consecutive trials across the three experimental blocks during the sham and otCS stimulation conditions. Error bars indicate the *SE*.

## 4 DISCUSSION

The current studied aimed to provide a proof of principle that the entrainment of ongoing neural oscillations by rhythmic visual stimulation can be replicated with cathodal otCS at the same frequency. To this end, we attempted to modulate the phase alpha oscillations in the posterior parietal cortex during a well-established visual detection task. Contrary to our hypothesis, there was no evidence that cathodal otCS stimulation pulses modulated target detection rates. We found that mean target detection rates during the otCS stimulation did not change as compared to sham stimulation. Furthermore, the sinusoidal pattern of the target detection rates did not seem to be strongly modulated by the cathodal otCS stimulation pulses compared to the sham. Together, these results did not provide significant evidence for 10 Hz cathodal otCS directly inducing modulation of alpha oscillations that can influence visual perception in a target detection task.

To the best of our analysis, cathodal otCS stimulation was not observed to modulate alpha oscillations and subsequent target detection rates. A major limitation of this study is that the efficacy of cathodal otCS can only by inferred from the perceptual and behavioural consequences of electrical stimulation during the target detection task. Although EEG was recorded throughout the experiment, we were not able to remove the otCS-induced artifacts. As a result, we have no direct electrophysiological evidence that the cathodal otCS stimulation interacted with the ongoing brain oscillations. Therefore, we cannot eliminate technical or methodological issues as the explanation for a lack of measurable behavior effects. For example, it is possible that the stimulation intensity or duration was not sufficient for inducing modulation of endogenous alpha oscillation. However, it is unlikely that stimulation intensity was too low to induce effects because previous studies have used similar intensities with observable effects (Moliadze et al., 2012; Neuling et al., 2015). Insufficient stimulation duration is also an unlikely explanation because there was no change in target detection rates compared to sham over course of the target detection task (see Figure 4). Furthermore, the three blocks of the target detection task took at least 10 mins which is considered enough time to induce effects in the ongoing oscillations (Antal et al., 2008; Thair et al., 2017).

It is also possible that using a 10 Hz stimulation frequency for all participants rather than matching the otCS frequency to each individuals’ peak alpha frequency reduced the efficacy of cathodal otCS. Several lines of evidence have shown that effective modulation of endogenous oscillations by periodic brain stimulation depends on matching the stimulation frequency to the rhythmic activity. For example, a study using optogenetic stimulation and multichannel slice electrophysiology found that a weak sine-wave electric field can enhance ongoing oscillatory activity, but only when the stimulation frequency was matched to the endogenous oscillation (Schmidt et al., 2014). Furthermore, a meta-analysis of fifty-one sham controlled experiments that investigated the effects of tACS on perception and cognitive performance, Schutter and Wischnewski (2016) found that performance is more likely to increase when tACS is administered based on individual spectral information. Together, these results suggest that the efficacy of cathodal otCS in the current study might have been greatly reduced because we did not control for inter-individual differences of endogenous alpha oscillations. However, using a 10 Hz stimulation frequency rather than matching the otCS frequency to individual peak frequencies might not have been as important a factor as it might seem. Specifically, even in the same participant, individual endogenous oscillatory activity varies during the course of a given task which could decrease the effects of stimulation even when the individual peak frequency was applied (Woods et al., 2016).

Another factor that could have reduced the efficacy of this method was that we did not control the timing of the otCS stimulation with regards to the target detection task. As a result, state-dependent differences in cortical activity across individuals prior to otCS may influence the effects of subsequent stimulation, introducing a possible source of variability (Silvanto and Pascual-Leone, 2008). However, this is an unlikely explanation because much of the variability due to differences across individuals would have been accounted for in the sham condition and by blocking on participants in the statistical analysis. Therefore, it is unlikely that state-dependent differences in cortical activity could significantly contribute to the lack of behavioral differences between the otCS and sham conditions in the target detection task.

In addition to the technical and methodological limitations mentioned above, individual differences in the brain’s susceptibility to otCS is another factor that may contribute to the lack of an observable effect. Anatomical variation including scalp-brain distance, gyral folding of the cerebral cortex, and thickness of corticospinal fluid layer and skull can have a significant impact on the effects of transcranial current stimulation (Nitsche et al., 2008; Opitz et al., 2015).

The results of the current study suggest that 10-Hz cathodal otCS stimulation does not directly induce modulation of alpha oscillations that can influence visual perception in a target detection task. Part of this null result might be explained by individual differences in peak alpha frequency, state-dependent changes in cortical activity, and susceptibility to otCS stimulation. However, technical and methodological issues might also contribute a lack of observable differences in visual perception. In the absence of electrophysiological evidence, it is important to be cautious about forming any firm conclusions based on the current study. Further research is needed to convincingly eliminate cathodal otCS stimulation as a means of modulating endogenous alpha oscillations in the posterior parietal area. However, the current study provides the first evidence supporting that conclusion.

## 6 AUTHOR CONTRIBUTIONS

SSS and KEM contributed conception and design of the study. SSS performed the statistical analysis. Both authors interpreted the data and wrote the article. Both authors approved the final version of the manuscript.

## 8 ACKNOWLEDGMENTS

The authors would like to thank Anna Kim for assistance with data collection.

